# Climate, human influence, and the distribution limits of the invasive European earwig, *Forficula auricularia*, in Australia

**DOI:** 10.1101/378570

**Authors:** Matthew P Hill, Matthew Binns, Paul A Umina, Ary A Hoffmann, Sarina Macfadyen

**Author notes:** **Corresponding author:** Matt Hill, CSIRO Data61, Black Mountain, ACT, Australia.

## Abstract

**BACKGROUND:** By modelling species-environment relationships of pest species, it is possible to understand potential limits to their distributions when they invade new regions, and their likely continued spread. The European earwig, *Forficula auricularia*, is a non-native invasive species in Australia that has been in the country for over 170 years. However, in the last few decades it has invaded new areas. Unlike in other countries, *F. auricularia* is a pest species of grains production in Australia. In this study we detail the Australian distribution of this species, adding new samples focussed around grain growing regions. Using this information we build global species distribution models for *F. auricularia* to better understand species-environment relationships.

**RESULTS:** Our models indicated that the distribution of *F. auricularia* is strongly associated with temperate through to semi-arid environments, a high winter rainfall and pronounced temperature seasonality. We identified regions that hold suitable, but as yet vacant, niche space for Australian populations, suggesting further potential for range expansion. Beyond climate, an index describing human influence on the landscape was important to understand the distribution limits of this pest. We identified regions where there was suitable climate space, but which *F. auricularia* has not occupied likely due to low levels of human impact.

**CONCLUSION:** Modelling the global distribution of a non-native pest species aided understanding of the regional distribution limits within Australia and highlighted the usefulness of human impact measures for modelling globally invasive insect species.

## 1 INTRODUCTION

The global costs of invasive insects are estimated at US$70 billion annually^1^, and new introductions continue to occur.^2^ Predicting the distribution, spread and potential impacts of non-native insects is an important, ongoing and challenging task.^3^ Effective management and policy formation require accurate predictions of potential distribution, especially for non-native economic pests, and species that are ecologically disruptive or impact human health.^4^ Building predictive models for mapping pest distributions not only helps to communicate risk to industry, but also increases understanding of ecological limits to species distributions and identification of processes that may promote invasiveness.

Climate suitability is a major predictor of establishment success and subsequent spread for insects, which has been exploited to produce distribution maps for use in management for close to 100 years (dating back to WC Cook^5–7^). The ability to translate species climatic or environmental tolerances into geographical prediction relates to the *niche*. If there is enough information on the direct physiological requirements of a species, obtained through experimental evidence and field-testing, it is possible to estimate components of the *fundamental niche* of a species (e.g. Kearney et al.^8^; Gutierrez et al.^9^). Presence/absence points for a species represent a physical expression of the niche, and theoretically approach the fundamental niche given complete sampling and no biotic interactions or constraints to dispersal. However, biotic interactions and barriers to movement are common and further restrict distributions, thus the *realised niche* describes the proportion of the fundamental niche that is expressed by a population.^10–12^ Nevertheless, for predicting establishment and potential distributions of many globally invasive insect species, the realised niche defined by climatic suitability is often a useful first approximation, unless there are substantial evolutionary shifts that facilitate niche change in an invaded region (e.g. Rey et al.^13^; Egizi et al.^14^).

There are several different approaches for characterising the niche and linking a species to its environmental requirements. Species distribution models (SDMs) (or ecological niche models) are the most commonly applied technique. These characterise species-environment relationships by correlating observations of species occurrence with covariates, typically in the form of interpolated climate or environmental indices generated through Geographical Information Systems (GIS). Species distribution models in this sense attempt to characterise something close to the realised niche. The accuracy of SDMs depends on all factors defining the niche boundaries, and hence distribution, being included at the appropriate modelled scale.^15^ This requirement means that choice of variables to include in the model and availability of occurrence records are both critical to producing meaningful predictions. While applying the same set of predictor variables across a range of species can give insight into general patterns of invasion,^16^ predicting the distributions of invasive species is not a ‘one size fits all’ approach.^17^ Predictor variables should ideally be chosen based on how directly they relate to a species’ physiology, and based on a low level of correlation between variables across ranges (see Petitpierre et al.^18^). Whilst climatic predictor variables are typically employed, responses to modified environments are also likely to contribute to invasion success of non-native insect species.^19,20^

Typically, the only occurrence data available are *ad hoc* observations of species presence, rather than rigorous and systematic sampling strategies that yield both presence and absence information. There will also be biases in occurrence records, depending on the status of the species in a region, and resources used for monitoring. For example, more observations are likely in regions where the species is a pest, however many species are not pests in their native range. Species distribution models constructed from the native range alone (with perhaps limited observations) may then be poor predictors of invasive ranges.^16, 21^ The inclusion of all known populations across the entire distribution can provide better characterisation of species-environment relationships. Additionally, a well-sampled species distribution across ranges allows SDMs to investigate which variables are relevant (proximal) in defining current distributions and to identify changes between native and invaded environments.^22^

The European earwig, *Forficula auricularia* L. (Dermaptera: Forficulidae) is a species with a broad native range extending across northern Africa, Europe and eastern Asia,^23^ with many non-native populations (e.g. United States of America and New Zealand), and introduced to Australia over 170 years ago.^24^ *Forficula auricularia* has been common throughout south-eastern Australia since the 1900s; despite only being first recorded in Western Australia 24 years ago, it is now common throughout south-western parts of this state.^24^ *Forficula auricularia* is often found in disturbed locations^23, 25^ and can be an important predator in apples,^26^ pear^27^ and kiwifruit^28^ orchards, but regarded as a plant-feeding pest in softer fruits such as stonefruit^29^, and sometimes as a contaminant during harvest.^30^ In Australia, *F. auricularia* is a pest in grains crops,^31, 32^ although internationally it is more typically considered a beneficial predator in grains systems.^33–36^ *Forficula auricularia* is generally found in locations with pronounced summer/winter seasonality. Populations can survive cold winters, where total annual rainfall exceeds 500 mm.^37^ The overall development of *F. auricularia* depends on temperature,^38^ with upper development temperature limits around 23-28°C.^38^ Temperatures above 24°C may reduce population size.^37^ The life history of *F. auricularia* is dependent on lineage; two reproductively isolated clades have been identified in Europe and North America, though they often coexist. Clade A produces one brood per year, while clade B can produce two broods per year.^25^ As far as we know, all Australian earwigs are from clade B.^24^ A better understanding of environmental factors that affect the spatial distribution of this species can help predict distribution limits in Australia, and aid targeted management strategies across local grains production areas.

In this study, we combine published records, field data, and pest reports to construct global species distribution models for *F. auricularia*. Using these models, we characterise species-environment relationships to predict the distribution of *F. auricularia* in Australia. We assess if the species occupies a different niche to populations elsewhere in the world, both native and introduced. Additionally, we test whether the species is in climatic equilibrium, whether it has expanded its niche, and whether there is potential for future range expansion. Given a putative association with human-impacted environments, we also assess how such impacts might change species-environment relationships and distributional limits, particularly with respect to Australian grains regions.

## 2 MATERIALS AND METHODS

### 2.1 Global spatial information

We collated georeferenced distribution records from various sources to construct a dataset of locations where *F. auricularia* has been recorded. The Global Biodiversity Information Facility (GBIF) is an open database that provides records from several sources to aid research. We used the R (version 3.4.4^39^) package ‘rgbif’^40^ to download all records for *F. auricularia*, which provided 26,012 records. The Atlas of Living Australia (ALA) is an open database for reports of any species found in Australia. We used the R package ‘ALA4R’^41^ to download occurrence data for *F. auricularia*, which gave 24 records. The Australian Pest Plant Database (APPD; Plant Health Australia 2001) is a closed database that contains records of pests and diseases of economically important plants in Australia. This yielded 238 records. All databases were accessed on 27.02.2018. In addition to these databases, we added further Australian data from a published field dataset,^24^ which yielded another 33 records. The data collected through field sampling and pest reporting services (up to 2014; see below) were included in this larger dataset to construct the SDMs.

We then examined the entire dataset and removed outliers to restrict the distribution to points that fell within Europe, North America, New Zealand and Australia (the first three comprising our “global” dataset). While *F. auricularia* is reported in Africa (as a native, except for South Africa where it is reported as invasive) and South America (Chile and Falkland Is.), there was too little information on its distribution within these regions. To address different resolutions of reported coordinates (and provide data at the same resolution as the predictor variables) we rescaled all observations to the grid cell level (10’, see below), which yielded 3,247 unique points.

### 2.2 Targeted field sampling in grains

To enhance the distribution dataset, we conducted targeted sampling of *F. auricula* over 2016-2017, specifically focussing on grain crops throughout Australia (see Supplementary Material 1.1 for map). Samples were taken in crops (canola, wheat, barley) using cardboard rolls that encourage earwigs to take refuge.^23,42^ The cardboard rolls consisted of single-sided corrugated cardboard 250 mm in width rolled to form 50 mm diameter cylinders with longitudinal corrugations. The rolls were inserted into a 200 mm length of 50 mm diameter polyvinyl chloride (PVC) pipe and were left out for seven days at each location. As earwigs are also ground-active,^43^ we used pitfall traps at the same sites.^44^ Each trap consisted of a PVC sleeve placed in the ground, flush with the soil surface. Vials 45 mm in diameter and 120 mL volume, containing 60 mL of 100% propylene glycol, were placed inside the sleeves and left open for seven days.

In addition to field samples, we obtained data from three Australian pest reporting services for the major grains growing regions of Australia. These reporting services collate incidences of pest outbreaks and reports from farmers, farm advisors, and other industry personnel in the respective regions (Western Australia, South Australia, New South Wales and Victoria). They involve PestFacts south-eastern (cesar; http://cesaraustralia.com/sustainable-agriculture/pestfacts-south-eastern), PestFacts South Australia (SARDI; http://www.pir.sa.gov.au/research/services/reports_and_newsletters/pestfacts_newsletter), and PestFax Western Australia (DAFWA; https://www.agric.wa.gov.au/newsletters/pestfax). These services yielded 17, seven and 53 localities with coordinates, respectively.

### 2.3 Environmental predictors and geographic extent

To characterise species-environment relationships of *F. auricularia* with broad scale variables, we employed the 19 bioclimatic variables from WorldClim 2.0^45^ at a 10’ resolution. These variables describe means, patterns and trends for temperature and precipitation observations in the interval 1970-2000. They were part of the ‘BIOCLIM’ package^46^ and are widely employed in species distribution modelling. The scale of 10’ (roughly 20 km^2^ at the equator) is also relevant to the model construction and transferability for broadly distributed species between large, distinct, geographical regions.^16,47,48^ We included a global aridity index from the CGIAR-CSI (Consortium for Spatial Information – Consultative Group for International Agriculture Research) database^49,50^ (http://www.cgiar-csi.org: accessed February 2018), as it correlates with the limit of the distribution of *Halotydeus destructor*, an invertebrate pest with a broadly similar Australian distribution.^22^ Within Australia, *F. auricularia* is rare in undisturbed habitats,^24^ so we also included a non-climatic environmental predictor that describes the impact of human influence (HII^51^) to examine how the global distribution may be associated with factors such as human population density, roads, agriculture and urban development. We consider this variable to indicate anthropogenic disturbance at a grid cell. The index ranges from 0 to 72, with higher scores indicating greater human influence.^51^ We also examined a soil classification layer, however this was not included in the modelling process but in examining model output (see Supplementary Material 2)

Prior to SDM and niche analyses, the geographic extent of the analysis was defined. The modelling extent, or background, is typically defined by restricting the geographic extent to environmental conditions similar to the environments held across the presence points, but that have not been occupied for a number of reasons. To achieve this, we selected the backgrounds for this study by using the Biome definitions of Olson et al.^52^ This approach has been applied successfully to other invasive species (e.g. Mateo et al.^53^; Hill et al.^16^) and provides a workable and repeatable background selection procedure. All biomes per continent that held a presence point were retained and used to create a single surface to extract background information.

As fewer variables are likely to result in models with better transferability, we examined which predictor may be more closely associated with the distribution of this species. We calibrated a Principal Components Analysis (PCA) across all available predictors for the entire study area, an effective method for choosing variables to aid transferability when the best predictive variables are unknown.^18^ Using R, we performed a PCA across all presence points to examine the variance and loadings of the different climatic predictor variables. This method identified aridity as explaining 99.8% of the total variance on the first axis, and seven bioclimatic variables on the second axis (temperature seasonality (bio04), annual precipitation (bio12), precipitation of the wettest month (bio13), precipitation of driest month (bio14), precipitation of wettest quarter (bio16), precipitation of driest quarter (bio17), bio18 = precipitation of warmest quarter (bio18), precipitation of coldest quarter (bio19)). We performed another PCA, this time removing aridity, to examine variance explained by the bioclimatic variables, and the first two axes of the PCA explained 84.4% and 13% of the total variance, respectively, with the same seven variables identified. While it was not important in terms of PCA loadings, the density of the HII across occurrences versus a null distribution indicated a clear positive association of *F. auricularia* with human impact, warranting its inclusion. We then generated 40,000 random points across the entire training background (global distribution) of *F. auricularia* and extracted predictor information for the eight variables. Correlations between the eight climatic variables were assessed using Kendall’s τ (tau) within the ‘corrplot’^54^ package in R before inclusion in the initial models (Supplementary Material 1.3).

### 2.4 Species distribution modelling

Our approach to species distribution modelling employed an iterative approach across five models (see Table 1). Firstly, a model was developed based on our predictor set of eight variables using default settings of Maxent as implemented through the ‘dismo’^55^ package in R. Maxent has been widely used for SDM purposes and provides a suitable framework for modelling when the data is presence-only (i.e. true absence data is not available – typical of insects^56^). Secondly, we changed the settings of Maxent to those of a point process model.^57^ While regression models examine the relationship between a random variable to covariates, point-process models work on the principal that the spatial location of the observed points is driven by the covariates, and investigate this by jointly modelling the location of the points with the expected intensity per unit area.^57^ An advantage over default Maxent settings here is that the predicted intensities from point-process models are scale invariant. Thirdly, we repeated the second model, but without the inclusion of the HII variable, to examine how a model with only climate variables performs. Following these initial three models, model selection was achieved through AICc (corrected Akaike Information Criterion) and BIC (Bayesian Information Criterion) using all data for training and the R package ‘rmaxent’^58^, and by examining AUC (Area under the curve of the receiver-operator characteristic) across 10 cross-validated replicates, using a 70:30 training:testing split.

**Table 1.** Model selection process. Columns give the variables used for each Maxent model: n (the number of occurrence records used for model training), k (the number of features with non-zero weights), ll (the negative log likelihood of the model), AIC (Akaike Information Criterion), AICc (sample size corrected AIC), and BIC (Bayesian Information Criterion) and AUC (Area Under Curve of the Receiver-Operator Characteristic).

| Model - settings | Variables | N | k | ll | AIC | AICc | BIC | AUC |
| --- | --- | --- | --- | --- | --- | --- | --- | --- |
| 1 – default | arid bio04 bio12 bio13<br>bio16 bio17 bio18<br>bio19 hii | 3113 | 156 | -<br>31858.68 | 64029.<br>35 | 64045.<br>92 | 64972.11 | 0.94 |
| 2 – point process | arid bio04 bio12 bio13<br>bio16 bio17 bio18<br>bio19 hii | 3113 | 52 | -<br>31445.68 | 62995.<br>35 | 62997.<br>15 | 63309.61 | 0.96 |
| 3 – point process | arid bio04 bio12 bio13<br>bio16 bio17 bio18<br>bio19 | 3272 | 221 | -<br>34412.12 | 69266.<br>24 | 69298.<br>41 | 70612.83 | 0.95 |
| 4 – point process | arid bio04 bio19 bio13<br>hii | 3113 | 164 | -<br>31381.26 | 63090.<br>53 | 63108.<br>89 | 64081.64 | 0.96 |
| 5 – point process | arid bio04 bio19 bio13 | 3272 | 163 | -<br>34998.17 | 70322.<br>34 | 70339.<br>55 | 71315.53 | 0.94 |

Variables: arid = Aridity index, bio04 = Temperature Seasonality (standard deviation \*100), bio12 = Annual Precipitation, bio13 = Precipitation of Wettest Month, bio14 = Precipitation of Driest Month, bio16 = Precipitation of Wettest Quarter, bio17 = Precipitation of Driest Quarter, bio18 = Precipitation of Warmest Quarter, bio19 = Precipitation of Coldest Quarter, hii = Human Influence Index (HII).

The variable contribution, permutation importance, and limiting factor analyses were then used to determine which variables contributed most, and where, in Australia. This led to a subset of predictor variables for subsequent analysis. We examined the modelled responses of the individual variables (percent contribution and permutation importance) and from these results (and examining the amount of correlation between variables (τ < 0.7)) removed an additional three variables. We then ran a fourth Maxent model on the reduced number of variables and compared this with the previous models, to examine if any important information had been discarded. Finally, for the fifth model, we again excluded the HII and compared this back to the other models to examine the influence of the non-climatic variable within the reduced predictor set.

### 2.5 Reciprocal distribution modelling

To examine how the geographical representation of the niche may differ between Australian and non-Australian populations of *F. auricularia*, we constructed SDMs based on subsets of the data (corresponding to different ranges), and reciprocally projected these models between ranges.^22, 48^ Here we aimed to determine how well the species-environment relationships of the global distribution can be characterised from the populations in Australia, and *vice versa*. To perform the reciprocal distribution models we used the same predictor variables and model settings as found through the SDM process, but only trained the model on a subset of the data, corresponding to either the ‘global’ (N. America, Europe, and New Zealand) or ‘Australia’ ranges. Once the models were constructed, we projected them to the reciprocal background: ‘Australia’ to ‘global’, ‘global’ to ‘Australia’. We then calculated a measure of niche overlap, Schoener’s D, and used the corresponding range point data to perform an AUC test.

### 2.6 Niche change analysis

To further understand how the niche of Australian populations may be different from the non-Australian populations in environmental space, we measured amounts of niche change as detailed elsewhere^47,16,59^, which involves a PCA across the combined backgrounds (global and Australia) and the respective ranges. All PCAs were rescaled to 100×100 cells and the densities of occurrence (the native and non-native ranges, respectively) were projected onto these surfaces. This procedure is undertaken to reduce bias from the different sizes of the two ranges and the relative amount of sampling undertaken. The two rescaled surfaces were then overlaid, and the amount of niche change measured. The three main measured components were the amount of overlap (again through Schoeners’ D), niche expansion (climates available in both ranges that have not been exploited in the native range) and niche unfilling (climates available in both ranges that have not been exploited in non-native range). We then examined these three components of niche change with and without the inclusion of HII, to test how much climate versus other abiotic factors (encapsulated in the HII) affected niche change and hence potential distribution.

## 3 RESULTS

### 3.1 Distribution information

The study contributed 39 new distribution localities from grains production systems in Australia through field surveys, adding to those from the pest reporting services to create a novel dataset of 100 *Forficula auricularia* distribution points (Supplementary Material 1.1). Nineteen sampled localities did not yield any *F. auricularia*. As these sites may indicate seasonal absence rather than complete absence, they were not included in our modelling.

### 3.2 Environmental variables and model selection

The predictor variables identified as being important and model selection criteria are shown in Table 1. Two models stand out as best performing: models 2 and 4. Model 2 has nine predictor variables, whereas model 4 has only five. The slight decrease in AUC and increase in AICc for selecting model 4 over 2 can be justified in the lower dimensionality, which is important to model transferability^60^ (i.e. the subsequent reciprocal distribution modelling and niche analysis use only subsets of the data and project across ranges). Our final predictor variable set, in order of importance, consisted of the human influence index (HII), bio04 (Temperature Seasonality (standard deviation *100), bio19 (precipitation of the coldest quarter), aridity index, and bio13 (precipitation of the wettest month). The variable response curves of model 4 are in Supplementary Material 1.4. As the aridity index is defined as mean annual precipitation divided by mean annual potential evapotranspiration, it represents precipitation availability over atmospheric water demand. Aridity values at around 0.2 (semi-arid) resulted in higher probabilities (for this variable), with arid (<0.1) and humid (> 0.65) locations being much less suitable. Beyond aridity, this predictor set indicates a strong association with winter rainfall (200-800 mm) and defined seasons (standard deviation of 3-4°C over 1970-2000, based on the standard deviation of monthly temperature averages) (Supplementary Material 1.4). Based on the PCA constructed on climatic information (Figure 1a), the Australian distribution is nested with the global distribution, and the distribution from the grains growing regions of Australia is nested within the Australian distribution (Supplementary Material 1.2). The density of occurrences across the HII layer (Figure 1b) shows that the global distribution of *F. auricularia* is associated with higher levels of human impact than expected at random. The modelled response for HII shows that higher values for this variable resulted in higher probabilities (Supplementary Material 1.4).

**Figure 1.**
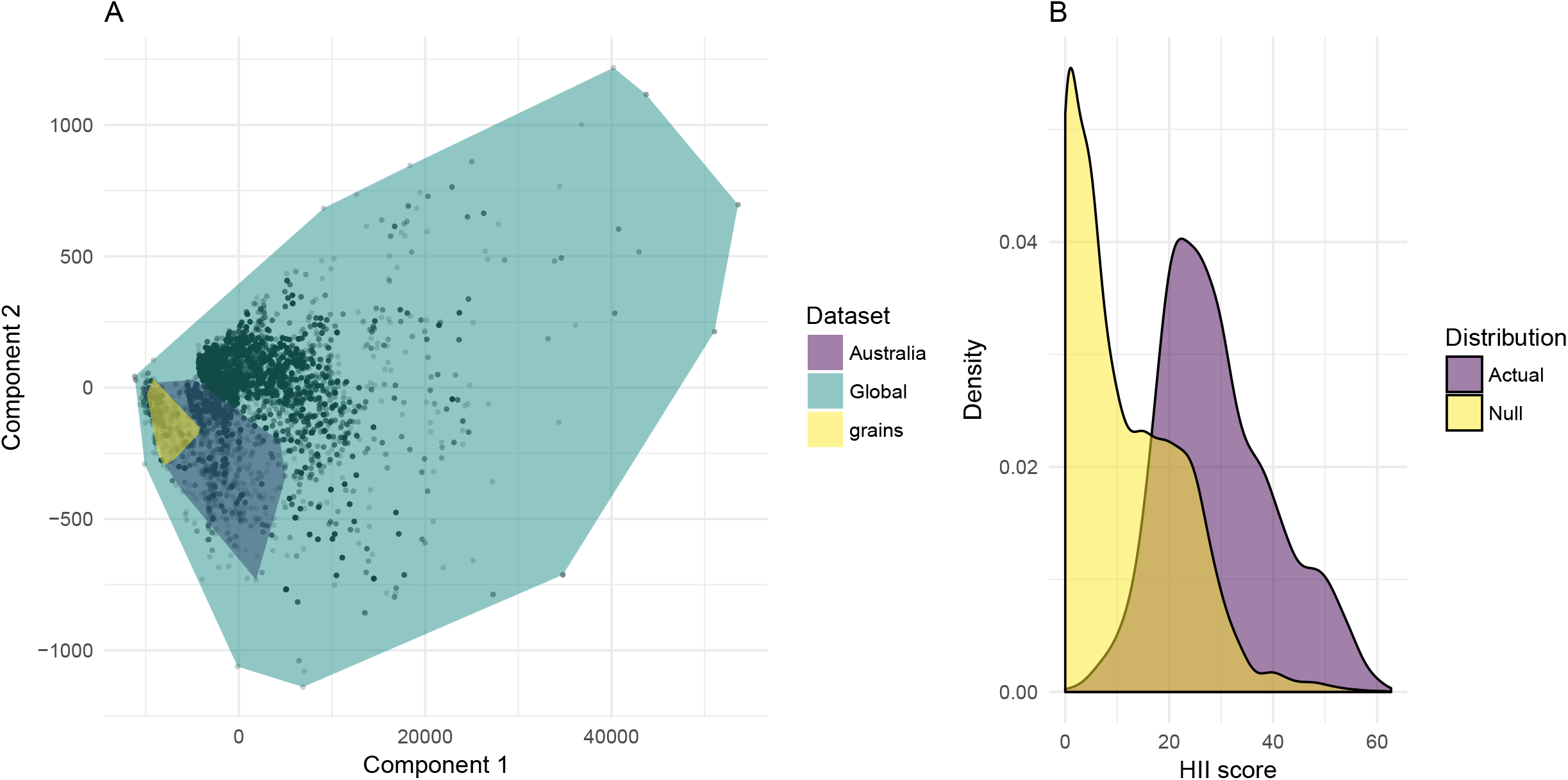
A) PCA of climatic variables showing the nesting of Australian data and Australian grains distribution data for *F. auricularia* within each other. Component 1 is the aridity index. Component 2 represents the other climatic variables used in the SDMs. B) Density of HII values for the global distribution of *F. auricularia* versus a null distribution of the same number of points at the grid cell level (n = 3113).

### 3.3 Model prediction

The model prediction in Australia shows habitat suitability for *F. auricularia* is mainly restricted to southern parts, corresponding to areas of strong seasonal differences in temperature, and a predominantly winter rainfall (Figure 2). These areas also correspond to the main grains cropping regions of southern Australia. In Western Australia, there is suitable habitat to the north of the currently observed distribution. The limiting factor analysis that accompanies this model prediction (Supplementary Material 1.5) shows that HII often restricts the distribution more than climatic variables, and aridity limits the distribution around the occurrence points. Areas such as south-west Tasmania and parts of eastern Victoria highlight regions where the climate is expected to be suitable (Supplementary Material 1.5), but perhaps these regions are not sufficiently disturbed to facilitate population establishment.

**Figure 2.**
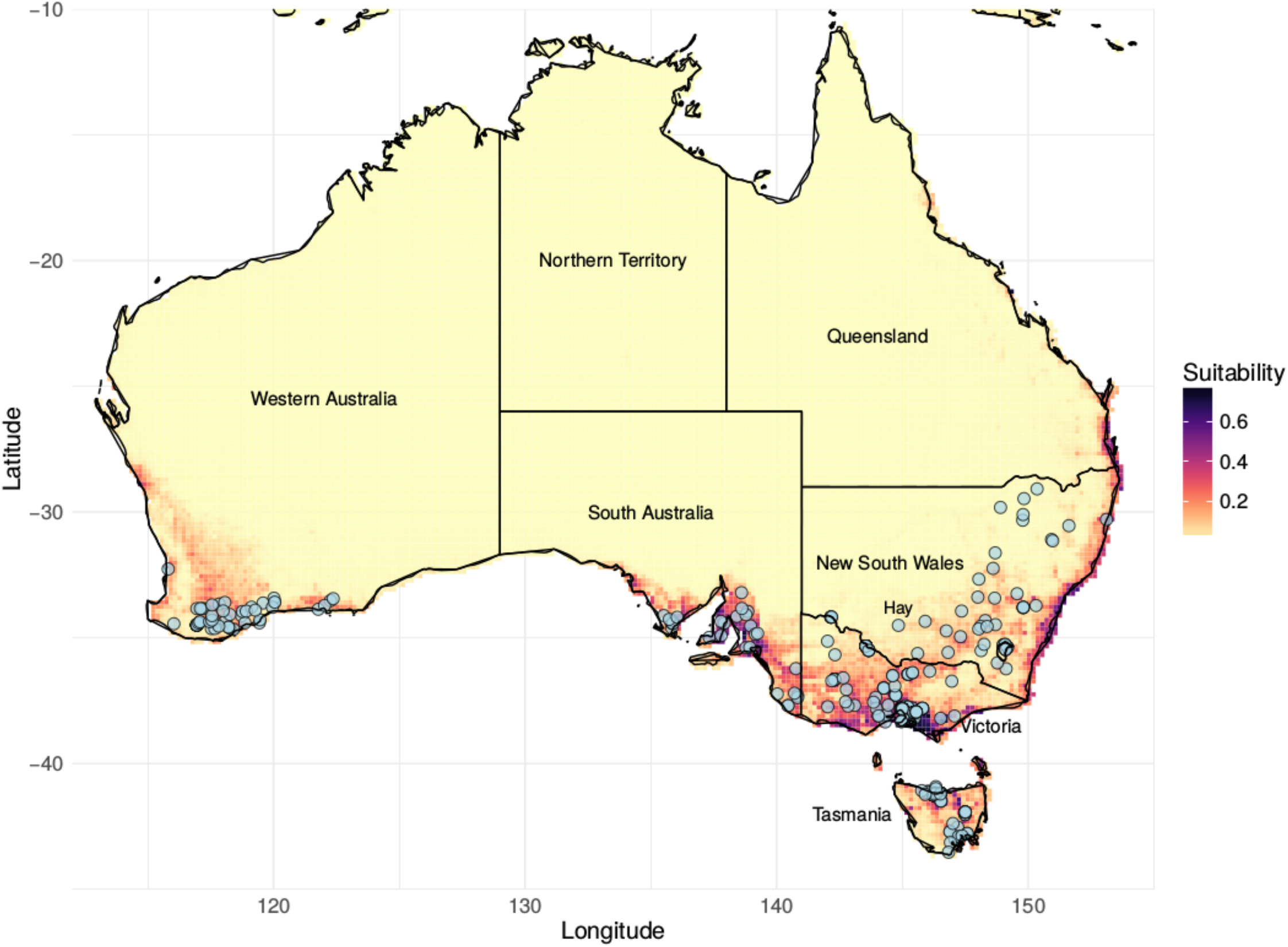
Modelled distribution of *Forficula auricularia* trained on all available data, predicted in Australia. This prediction is the final model selected for optimal settings and variables (see text). The scale ranges from 0-1, with 1 indicating high suitability, and 0 indicating not suitable. Light blue points indicate localities where *F. auricularia* has been found.

### 3.4 Reciprocal distribution modelling

For the reciprocal distribution modelling (RDM) (Figure 3), the global model projected globally is the predicted distribution od *F. auricularia* given our dataset (Figure 3a). This global model projected well to the Australian background with an AUC of 0.83 and a Schoener’s *D* score of 0.89 (Figure 3b), indicating high transferability and overlap in geographical space. The Australian model projected to the global background with an AUC of 0.94 and a Schoener’s *D* score of 0.82 (Figure 3c). These overlap scores indicate that there is little change in niche dimensions between the global and Australian distributions. The Australian-trained model does give a much denser prediction for Australia than the globally-trained one, indicating that the species-environment relationships are characterised slightly differently on this data subset (Figure 3d).

**Figure 3.**
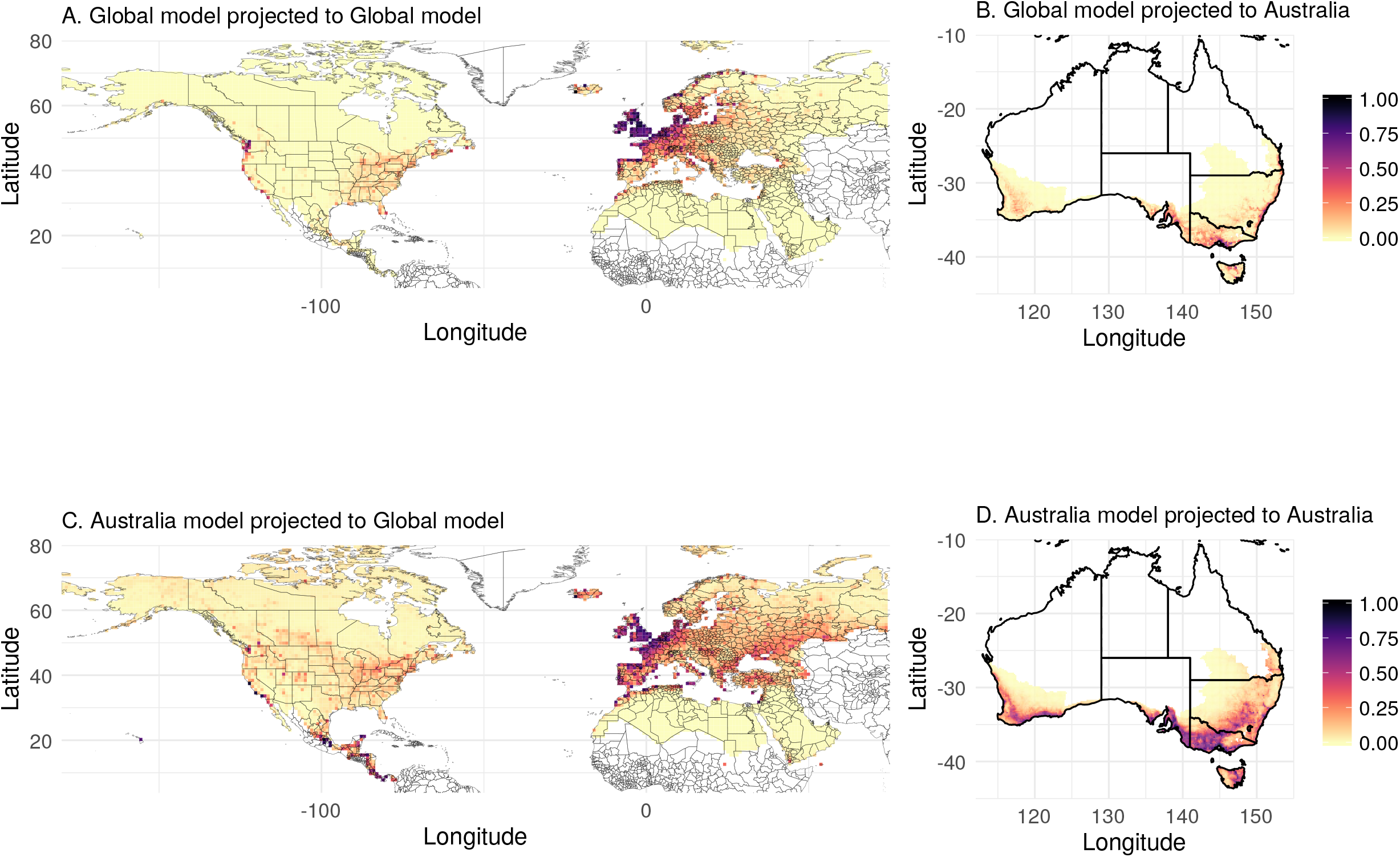
Reciprocal Distribution Modelling of *F. auricularia* between the ‘global’ distribution, and Australia. For mapping purposes, only the Northern Hemisphere is shown; the New Zealand data was, however, included in these analyses. The scale ranges from 0-1, with 1 indicating high suitability, and 0 indicating not suitable. The top two panels (A & B) show the projections made from a model trained only on the data from outside Australia. A) The projection into the ‘global’ range. B) The projection into Australia. The bottom two panels (C & D) are the projections made from a model trained only on the data from Australia. C) The projection from Australia to the ‘global’ range, D) The projection to Australia. The overlap (Schoeners’ *D*) between panels A & C, *D* = 0.9 and the overlap between panels B & D, *D* = 0.84.

### 3.5 Niche change

The niche change analysis shows low amounts of overlap in the environmental space occupied by the Australian and global distributions (D = 0.16), suggesting the global distribution is broader, and the Australian distribution occupies a subset of the total range of environments of *F. auricularia* (Figure 4). There is no evidence for niche expansion in Australia, again suggesting the Australian populations occupy a subset of the global niche, rather than any new environments (Table 2). Both niche change analyses also show high amounts of unfilling, which suggests that *F. auricularia* is not currently found in all possible environments in Australia) Table 2). Importantly, the niche change score for unfilling was much lower when the HII was included. In summary, the reciprocal distribution modelling and niche change analysis demonstrate that the Australian distribution falls within the environmental ranges expected from the global distribution (Figure 4, Table 2).

**Figure 4.**
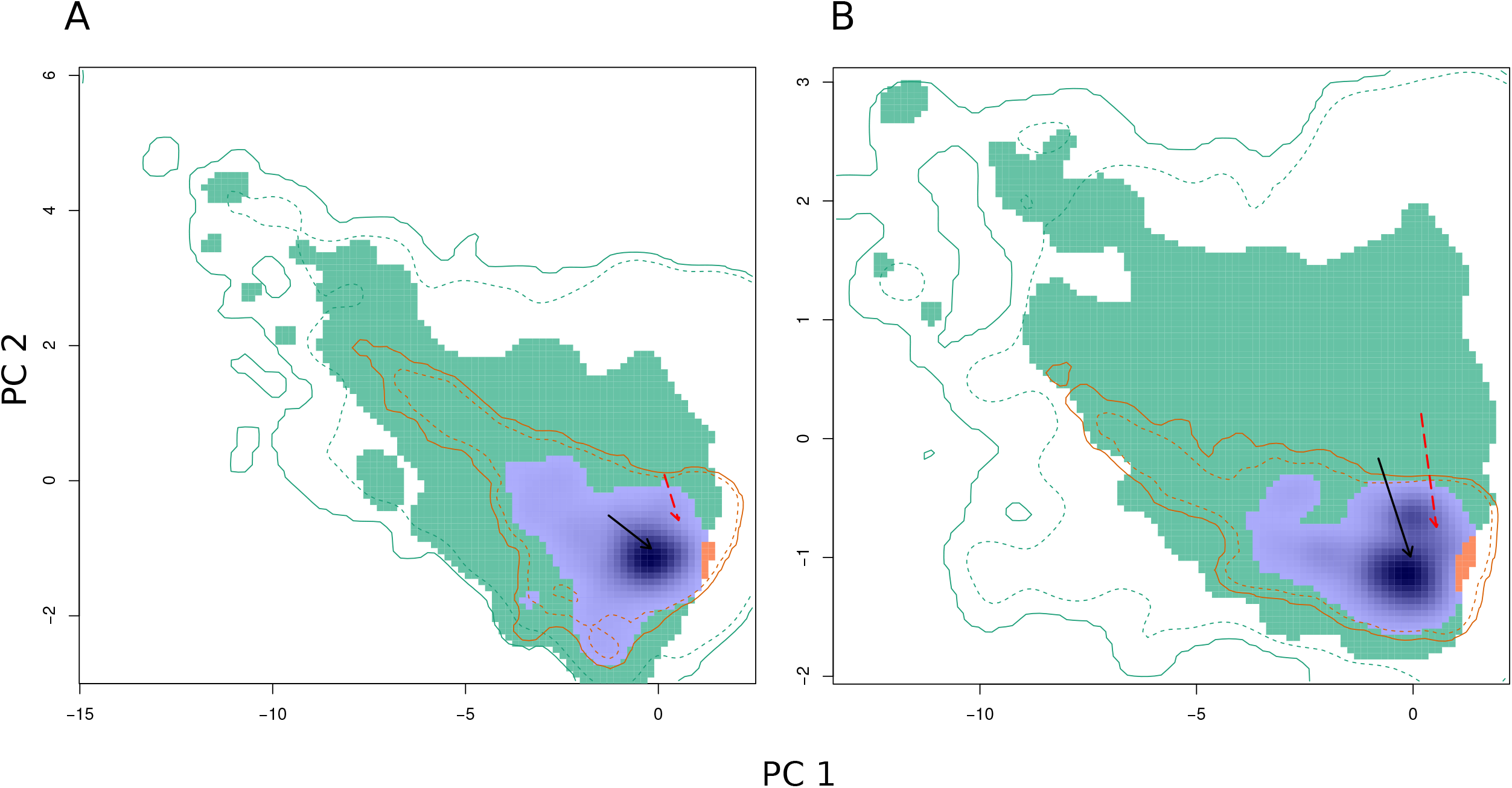
Niche change analysis for *Forcifula auricularia* in Australia compared to its global distribution. The green region indicates the amount of niche unfilling, the red the amount of niche expansion, and the blue the niche stability. The solid red line indicates the climates available in the non-native range (Australia) and the dashed line represents the climates at the 75^th^ percentile. The sold green line indicates the climates available in the native (global) range. A) Final climate dataset with the HII included. B) Final climate dataset without HII. See Table 2 for scores of the different niche change components.

**Table 2.**
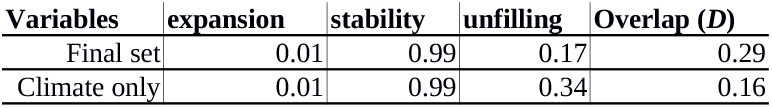
Niche change metrics for populations *Forficula auricularia* in Australia versus the ‘global’ distribution (see Figure 4 for descriptions). The overlap is Schoeners’*D*.

## 4 DISCUSSION AND CONCLUSIONS

*Forficula auricularia* is widely distributed across southern Australia. By using a comprehensive global dataset to characterise species-environment relationships and examining niche dynamics, we provide evidence that the distribution limits are not met and there is potential for future spread. *Forficula auricularia* displays relationships with climate that align with current knowledge of its physiology, and result in spatial overlap with the major grains growing regions of Australia. Further, *F. auricularia* has a strong association with human influenced habitat, an obvious attribute of broad-acre agricultural landscapes. Whether this is related to the pest status in Australian grains remains an interesting question, however the association of *F. auricularia* with human influenced habitat appears linked to the species’ ability to occupy certain environments, as we identified regions where there was suitable climate space, but which *F. auricularia* has not occupied likely due to a low level of human impact.

Aridity is the most important variable in limiting the distribution of *F. auricularia*. As for many invertebrate species, humidity is critical to prevent desiccation of *F. auricularia* eggs and nymphs.^38^ The other important predictors identified, winter rainfall and temperature seasonality, are typical attributes of the main grains growing regions of southern Australia. Winter rainfall is a critical component to successful grains cropping, supporting the initial growth of several economically important crops that are then typically harvested in spring and summer. Temperature seasonality further reflects components of the life-cycle. *Forficula auricularia* is likely to occupy below-ground nests when crops are sown (winter), and then emerge around the same time the crop is maturing (late spring-summer). Our study highlights the importance of using covariates beyond climate; in particular, human-impacted environments seem important for a number of non-native invasive insect species.^16^ Human influence may contribute to the distribution of *F. auricularia*, either directly, such as by reducing the heterogeneity of habitat (e.g. biotic homogenization through agricultural monocultures) and providing new host plants, or indirectly by reducing species that may compete with or predate.^61, 62^

For a few regions in Australia our models predict suitable habitat where *F. auricularia* has not been observed. The largest of these is in Western Australia (WA) to the north of current observations (Figure 2). Interestingly this was found to be suitable regardless of whether the human influence index was included. Quarrell et al.^24^ speculated that inland populations at Hay in New South Wales (Figure 2) (where our models predicted marginal suitability) are indicative of a species that can exist in a variety of environments, and therefore able to spread into new regions like those in WA. As *F. auricularia* has only been in WA for 24 years it may still be in the process of spreading to its limits (see Quarrell et al.^24^). Alternatively, there could be soil properties, or other abiotic components not included here limiting its distribution. However, we found no evidence for soil-related effects in that region (Supplementary Material 2). Finally, our models are calibrated on a global dataset that encompasses diverse populations across a broad range of climates; while this helps determine the species-environment relationships, it may identify regions suitable for a species as a whole, but not for a local population. Quarrell et al.^24^ found that, unlike for the rest of Australia, populations in WA were started by only a few individuals, perhaps limiting adaptive capacity and the ability of this population to expand further.

The broad distribution of *F. auricularia* across a range of agricultural environments suggests that the species either has a generalist nature, or that it has undergone adaptive shifts to allow it to move onto new hosts and landscapes, including grains, during its invasion.^63^ Some invasive insects, for example *Wasmannia auropunctata* and *Leptinotarsa decemlineata*, likely adapted to anthropogenic landscape changes in their home ranges (forests replaced with plantations, provision of new agricultural hosts) before becoming highly invasive species.^64^ Other species, particularly agricultural pests like *Ostrinia nubilalis* and *Daktulosphaira vitifoliae*, have undergone ‘bridgehead’ scenarios, enabling a single evolutionary shift towards invasiveness in an intermediate invaded region to then facilitate subsequent invasions.^63^ Given the strong association of *F. auricularia* with human-impacted environments, comparisons of *F. auricularia* populations from different landscapes should provide insights into this issue.

To better understand species-environment relationships in *F. auricularia*, additional occurrence information from places like South Africa, Chile, and the Middle East is needed. Consistent reporting across ranges is an issue with many globally invasive species,^65^ as surveying and reporting efforts between countries vary due to regional impact of the pest, reporting incentive, and resources available.^66^ Reporting bias is further complicated by the fact that *F. auricularia* is not considered a pest in some agricultural contexts. While the scale of predictors used in this study is suited to developing a range of hypotheses, some of the fine-scale behaviour and interactions with other species and environmental parameters could not be tested. For instance, predators such as ants or spiders as well as competitors may limit the distribution of this species. Other earwigs are found throughout the grain growing regions of Australia,^67,68^ although their abundance and pest/beneficial status is not understood. These include from the family Anisolabididae and the genera *Labidura* and Nala.^32,68^ To our knowledge, *F. auricularia* and *Nala lividipes* are the only confirmed non-native species in these systems, although other species (e.g.cryptic Anisolabididae) may have been introduced. Further work is required to understand the distributional limits of these other earwig species, along with potential niche overlap and competitive interactions.

We have shown that species-environment relationships characterized using SDMs provide insight into potential factors that limit distributions. While it was possible to map the potential distribution of *F. auricularia*, predictions around population abundance and pest status require empirical research and different models. Mechanistic models, including temperature-based day-degree models, can help predict timing and emergence of *F. auricularia* in grains systems. Further, while the human influence index affected the distribution of *F*. a*uricularia*, additional work is required to understand the components of human impact critical to the distribution of *F. auricularia*. This information in turn may indicate whether the species is an opportunistic pest of grain crops, or if grain production systems have facilitated its spread.

## ACKNOWLEDGEMENTS

We acknowledge Michael Nash, Joanne Holloway, Maarten Van Helden, Thomas Heddle and Dusty Severtson for a subset of earwig distribution points from Australia used in this study. Justine Murray and Madeleine Barton provided valuable comments on a draft of this manuscript. Thanks to SARDI, DAFWA and cesar for kindly providing historical pest reporting records. This work was supported by the Grains Research & Development Corporation (project CSE00059).

## SUPPORTING INFORMATION

Supporting information may be found in the online version of this article

Dataset of new Australian occurrences and R code for SDM and niche analyses is available at figshare: 10.6084/m9.figshare.6865031

## REFERENCES

1. Bradshaw CJ, Leroy B, Bellard C, Roiz D, Albert C, Fournier A, Barbet-Massin M, Salles JM, Simard F, Courchamp F. Massive yet grossly underestimated global costs of invasive insects. Nat Commun 7:12986 (2016).

2. Seebens H, Blackburn TM, Dyer EE, Genovesi P, Hulme PE, Jeschke JM, Pagad S, Pyšek P, Winter M, Arianoutsou M, Bacher S. No saturation in the accumulation of alien species worldwide. Nat Commun 8:14435 (2017).

3. Hill MP, Clusella-Trullas S, Terblanche JS, Richardson DM. Drivers, impacts, mechanisms and adaptation in insect invasions. Biol Invasions 18:883–891 (2016).

4. Venette RC, Kriticos DJ, Magarey RD, Koch FH, Baker RH, Worner SP, Gómez Raboteaux NN, McKenney DW, Dobesberger EJ, Yemshanov D, De Barro PJ. Pest risk maps for invasive alien species: a roadmap for improvement. BioScience 60:349–62 (2010).

5. Cook WC. The distribution of the pale western cutworm, *Porosagrotis orthogonia* Morr.: A study in physical ecology. Ecology 5:60–9 (1924).

6. Cook WC. A bioclimatic zonation for studying the economic distribution of injurious insects. Ecology 10:282–93 (1929).

7. Cook WC. Notes on predicting the probable future distribution of introduced insects. Ecology 12:245–7 (1931).

8. Kearney M, Porter W. Mechanistic niche modelling: combining physiological and spatial data to predict species’ ranges. Ecol Lett 12:334–50 (2009).

9. Gutierrez AP, Ponti L, Cooper ML, Gilioli G, Baumgärtner J, Duso C. Prospective analysis of the invasive potential of the European grapevine moth *Lobesia botrana* (Den. & Schiff.) in California. Agric For Entomol 14:225–38 (2012).

10. Austin MP. Spatial prediction of species distribution: an interface between ecological theory and statistical modelling. Ecol Modell 157:101–18 (2002).

11. Soberón J. Grinnellian and Eltonian niches and geographic distributions of species. Ecol Lett 10:1115–23 (2007).

12. Colwell RK, Rangel TF. Hutchinson’s duality: the once and future niche. Proc Natl Acad Sci USA 106:19651–8 (2009).

13. Rey O, Estoup A, Vonshak M, Loiseau A, Blanchet S, Calcaterra L, Chifflet L, Rossi JP, Kergoat GJ, Foucaud J, Orivel J. Where do adaptive shifts occur during invasion? A multidisciplinary approach to unravelling cold adaptation in a tropical ant species invading the Mediterranean area. Ecol Lett 15:1266–75 (2012).

14. Egizi A, Fefferman NH, Fonseca DM. Evidence that implicit assumptions of ‘no evolution’of disease vectors in changing environments can be violated on a rapid timescale. Phil Trans R Soc B 370:20140136 (2015).

15. Pearson RG, Dawson TP. Predicting the impacts of climate change on the distribution of species: are bioclimate envelope models useful?. Glob Ecol Biogeog 12:361–71 (2003).

16. Hill MP, Gallardo B, Terblanche JS. A global assessment of climatic niche shifts and human influence in insect invasions. Glob Ecol Biogeog 26:679–89 (2017).

17. Sutherst RW. Pest species distribution modelling: origins and lessons from history. Biol Invasions 16:239–56 (2014).

18. Petitpierre B, Broennimann O, Kueffer C, Daehler C, Guisan A. Selecting predictors to maximize the transferability of species distribution models: lessons from cross continental plant invasions. Glob Ecol Biogeog 26:275–87 (2017).

19. Roura-Pascual N, Hui C, Ikeda T, Leday G, Richardson DM, Carpintero S, Espadaler X, Gómez C, Guénard B, Hartley S, Krushelnycky P. Relative roles of climatic suitability and anthropogenic influence in determining the pattern of spread in a global invader. Proc Natl Acad Sci USA 18:201011723 (2010).

20. Magagula CN, Monadjem A, Dlamini WM. Predicted regional and national distribution of *Bactrocera dorsalis* (syn. *B. invadens*)(Diptera: Tephritidae) in southern Africa and implications for its management. Afr Entomol 23:427–37 (2015).

21. Broennimann O, Guisan A. Predicting current and future biological invasions: both native and invaded ranges matter. Biology Lett 4:585–9 (2008).

22. Hill MP, Hoffmann AA, Macfadyen S, Umina PA, Elith J. Understanding niche shifts: using current and historical data to model the invasive redlegged earth mite, *Halotydeus destructor*. Divers Distrib 18:191–203 (2012).

23. Lamb RJ, Wellington WG. Life history and population characteristics of the European earwig, *Forficula auricularia* (Dermaptera: Forficulidae), at Vancouver, British Columbia. Can Entomol 107:819–24 (1975).

24. Quarrell SR, Arabi J, Suwalski A, Veuille M, Wirth T, Allen GR. The invasion biology of the invasive earwig, *Forficula auricularia* in Australasian ecosystems. Biol Invasions 20:1553–65 (2018).

25. Guillet S, Guiller A, Deunff J, Vancassel M. Analysis of a contact zone in the *Forficula auricularia* L.(Dermaptera: Forficulidae) species complex in the Pyrenean Mountains. Heredity. 85:444 (2000).

26. Suckling DM, Burnip GM, Hackett J, Daly JC. Frass sampling and baiting indicate European earwig (*Forficula auricularia*) foraging in orchards. J Appl Entomol 130:263–7 (2006).

27. Moerkens R, Leirs H, Peusens G, Gobin B. Are populations of European earwigs, *Forficula auricularia*, density dependent? Entomol Exp Appl 130:198–206 (2009).

28. Logan DP, Maher BJ, Rowe CA. Predation of diaspidid scale insects on kiwifruit vines by European earwigs, *Forficula auricularia*, and steel-blue ladybirds, *Halmus chalybeus*. BioControl 62:469–79 (2017).

29. Bower CC. Control of European earwig, ‘*Forficula auricularia*’ L., in stone fruit orchards at Young, New South Wales. The Journal of the Entomological Society of New South Wales 24:11 (1992).

30. Gordon SC, Cormack MR, Hackett CA. Arthropod contamination of red raspberry (*Rubus idaeus* L.) harvested by machine in Scotland. J Hortic Sci 72:677–85 (1997).

31. Gu H, Fitt GP, Baker GH. Invertebrate pests of canola and their management in Australia: a review. Aust J Entomol 46:231–43 (2007).

32. Murray DA, Clarke MB, Ronning DA. Estimating invertebrate pest losses in six major A ustralian grain crops. Aust J Entomol 52:227–41 (2013).

33. Sunderland KD, Vickerman GP. Aphid feeding by some polyphagous predators in relation to aphid density in cereal fields. J Appl Ecol 17:389–96 (1980).

34. Sunderland KD, Crook NE, Stacey DL, Fuller BJ. A study of feeding by polyphagous predators on cereal aphids using ELISA and gut dissection. J Appl Ecol 24:907–33 (1987).

35. Manyuli T, Kyamanywa S, Luther GC. Capability of *Forficula auricularia* Linnaeus (Dermaptera: Forficulidae) to prey on *Aphis craccivora* Koch (Homoptera: Aphididae) in eastern and Central Africa. Agronomie Africaine 20:49–58 (2008).

36. Corpuz MR, Raymundo PP. Integrated pest management strategies to sustain corn productivity. Commun Agric Appl Biol Sci 75:411–6 (2010).

37. Crumb SE, Eide PM, Bonn AE. The European earwig. US Department of Agriculture (1941).

38. Moerkens R, Gobin B, Peusens G, Helsen H, Hilton R, Dib H, Suckling DM, Leirs H. Optimizing biocontrol using phenological day degree models: the European earwig in pipfruit orchards. Agric For Entomol 13:301–12 (2011).

39. R Core Team. R: A Language and Environment for Statistical Computing, R Foundation for Statistical Computing, Vienna, Austria. (2018).

40. Chamberlain S, Ram K, Barve V, Mcglinn D, Chamberlain MS. rgbif: Interface to the Global ‘Biodiversity’ Information Facility API. R package version 0.9.9. https://CRAN.Rproject.org/package=rgbif (2017).

41. Raymond B, VanDerWal J, Belbin L. ALA4R: Atlas of Living Australia (ALA) Data and Resources in R. R package version 1.6.0. https://CRAN.R-project.org/package=ALA4R (2018).

42. Quarrell SR. The chemical ecology, genetics and impact of the European earwig in apple and cherry orchards (Doctoral dissertation, University of Tasmania) (2013).

43. Frank SD, Wratten SD, Sandhu HS, Shrewsbury PM. Video analysis to determine how habitat strata affects predator diversity and predation of *Epiphyas postvittana* (Lepidoptera: Tortricidae) in a vineyard. Biol Control 41:230–6 (2007).

44. Epstein DL, Zack RS, Brunner JF, Gut L. Brown JJ. Effects of broad-spectrum insecticides on epigeal arthropod biodiversity in Pacific Northwest apple orchards. Environ Entomol 29:340–348 (2000).

45. Fick, SE, Hijmans RJ. WorldClim 2: new 1km spatial resolution climate surfaces for global land areas. Int J Climatol. 37:4302–4315 (2017).

46. Busby J. BIOCLIM-a bioclimate analysis and prediction system. Plant Protection Quarterly 6:8–9 (1991).

47. Broennimann O, Fitzpatrick MC, Pearman PB, Petitpierre B, Pellissier L, Yoccoz NG, Thuiller W, Fortin MJ, Randin C, Zimmermann NE, Graham CH. Measuring ecological niche overlap from occurrence and spatial environmental data. Glob Ecol Biogeog 21:481–97 (2012).

48. Fraimout A, Monnet AC. Accounting for intraspecific variation to quantify niche dynamics along the invasion routes of *Drosophila suzukii*. Biol Invasions DOI:10.1007/s10530–018–1750-z (2018).

49. Zomer RJ, Bossio DA, Trabucco A, Yuanjie L, Gupta DC, Singh VP. Trees and water: smallholder agroforestry on irrigated lands in Northern India. IWMI, Colombo Sri Lanka (2007).

50. Zomer RJ, Trabucco A, Bossio DA, Verchot LV. Climate change mitigation: A spatial analysis of global land suitability for clean development mechanism afforestation and reforestation. Agric Ecosyst Environ 126:67–80 (2008).

51. Sanderson EW, Jaiteh M, Levy MA, Redford KH, Wannebo AV, Woolmer G. The human footprint and the last of the wild: the human footprint is a global map of human influence on the land surface, which suggests that human beings are stewards of nature, whether we like it or not. Bioscience 52:891–904 (2002).

52. Olson DM, Dinerstein E, Wikramanayake ED, Burgess ND, Powell GV, Underwood EC, D’amico JA, Itoua I, Strand HE, Morrison JC, Loucks CJ. Terrestrial Ecoregions of the World: A New Map of Life on Earth. BioScience. 51:933–8 (2001).

53. Mateo RG, Broennimann O, Petitpierre B, Muñoz J, van Rooy J, Laenen B, Guisan A, Vanderpoorten A. What is the potential of spread in invasive bryophytes? Ecography 38:480–7 (2015).

54. Wei T, Simko V. R package “corrplot”: visualization of a correlation matrix (Version 0.84). Available from https://github.com/taiyun/corrplot (2017).

55. Hijmans RJ, Phillips S, Leathwick J, Elith J. R Package ‘dismo’: Species distribution modeling. R package version 1.1–4. https://CRAN.R-project.org/package=dismo (2017).

56. Lobo JM. The use of occurrence data to predict the effects of climate change on insects. Curr Opin Insect Sci 17:62–8 (2016).

57. Renner IW, Elith J, Baddeley A, Fithian W, Hastie T, Phillips SJ, Popovic G, Warton DI. Point process models for presence only analysis. Methods Ecol Evol 6:366–79 (2015).

58. Baumgartner J, Wilson P. rmaxent: Tools for working with Maxent in R. R package version 0.7.11.9000. Available from: https://github.com/johnbaums/rmaxent (2018).

59. Guisan A, Petitpierre B, Broennimann O, Daehler C, Kueffer C. Unifying niche shift studies: insights from biological invasions. Trends Ecol Evol 29:260–269 (2014).

60. Jiménez-Valverde A, Peterson AT, Soberón J, Overton JM, Aragón P, Lobo JM. Use of niche models in invasive species risk assessments. Biol Invasions 13:2785–97 (2011).

61. McKinney ML, Lockwood JL. Biotic homogenization: a few winners replacing many losers in the next mass extinction. Trends Ecol Evol 4:450–3 (1999).

62. Hendrickx F, Maelfait JP, Van Wingerden W, Schweiger O, Speelmans M, Aviron S, Augenstein I, Billeter R, Bailey D, Bukacek R, Burel F. How landscape structure, land use intensity and habitat diversity affect components of total arthropod diversity in agricultural landscapes. J Appl Ecol 44:340–51 (2007).

63. Guillemaud T, Ciosi M, Lombaert E, Estoup A. Biological invasions in agricultural settings: insights from evolutionary biology and population genetics. C R Biol 334:237–46 (2011).

64. Hufbauer RA, Facon B, Ravigne V, Turgeon J, Foucaud J, Lee CE, Rey O, Estoup A. Anthropogenically induced adaptation to invade (AIAI): contemporary adaptation to human altered habitats within the native range can promote invasions. Evol Appl 5:89–101 (2012).

65. Mang T, Essl F, Moser D, Karrer G, Kleinbauer I, Dullinger S. Accounting for imperfect observation and estimating true species distributions in modelling biological invasions. Ecography 40:1187–97 (2017).

66. Bebber DP, Holmes T, Smith D, Gurr SJ. Economic and physical determinants of the global distributions of crop pests and pathogens. New Phytol 202:901–10 (2014).

67. Allsopp PG, Radford BJ. Sowing techniques for sorghum on soil infested with black field earwigs and sugarcane wireworms. Queensland Journal of Agricultural and Animal Sciences (Australia) 44:15–19(1987).

68. Simpson GB, Mayer DG. Morphometric analysis of variation in *Nala lividipes* (Dufour) and *Labidura truncata* Kirby (Dermaptera: Labiduridae). Aust J Entomol 29:287–94 (1990).

